# Rise and Fall of a Superbug: Analyzing the epidemic success of a distinct clone of vancomycin-resistant *Enterococcus faecium* ST117/CT71 in Germany, 2017 - 2024

**DOI:** 10.64898/2026.11.10.765572

**Authors:** Robert E. Weber, Guido Werner, Jennifer K. Bender, Berit Haldemann, Oliver Drechsel, Martin A. Fischer

## Abstract

Vancomycin-resistant *Enterococcus faecium* (VR*Efm*) is a major cause of healthcare-associated infections, yet outbreak analysis remains challenging, particularly for prevalent lineages. One of the most striking examples is the Sequence Type (ST) 117/Complex Type (CT) 71/*vanB* lineage, whose rapid expansion across Germany coincided with a sharp rise in VRE prevalence observed around 2019. Defining reliable genomic thresholds remains particularly difficult, as current core-genome Multi-Locus Sequence Typing (cgMLST) cutoffs are based largely on expert consensus or limited case series.

Here, we evaluate the discriminatory performance of cgMLST and split k-mer analysis (SKA) for resolving relatedness within ST117/CT71 and investigate whether low core-genome diversity represents a lineage-specific trait or a population-wide property of *E. faecium*. While cgMLST captures the overall population structure of ST117/CT71, SKA provided higher resolution, distinguishing isolates identical by cgMLST. Analysis of 2,505 genomes showed that ST117/CT71 does not exhibit reduced allelic variability compared with other hospital-adapted lineages, suggesting that limited core-genome diversity is a broader population characteristic. Complementary transposon analyses further identified two temporally distinct ST117/CT71 sublineages carrying different Tn1549-like structures.

To assess allelic variation within confirmed transmission events, we analyzed 138 outbreak clusters from 96 hospitals. Outbreak-associated isolates showed sharply compressed allelic differences (median = 1), with 90% of pairwise comparisons falling within ≤9 alleles. Thus, the commonly applied ≤20-allele threshold risks overestimating genomic relatedness and grouping unrelated isolates.

Based on these data, we propose lowering the cgMLST alert threshold to ≤9 alleles for initial cluster detection of local outbreaks, followed by confirmatory high-resolution analyses such as SKA.

**Importance:** Defining reliable genomic thresholds for outbreak cluster detection remains a central challenge in bacterial genomic surveillance. Current cgMLST thresholds, largely based on expert consensus or limited case series, may overestimate genetic relatedness. Using VR*Efm* as an exemplar organism, we demonstrate that constrained core-genome diversity is a population-wide characteristic of clade A1 lineages rather than specific to ST117/CT71, limiting the discriminatory power of cgMLST approaches. Combining population-wide allelic comparisons with hospital outbreak data, we propose a data-driven cgMLST alert threshold of ≤9 alleles, followed by high-resolution analysis to improve outbreak detection. Furthermore, we show that the nationwide expansion of two closely related *vanB*-carrying ST117/CT71 sublineages coincided with the sharp rise in German VRE prevalence and incidence around 2019. These findings highlight how microevolutionary changes within dominant lineages can shape national epidemiological trends while remaining obscured by conventional typing approaches.

## Introduction

*Enterococcus faecium* (*E. faecium*) is a major cause of hospital-acquired infections, particularly bloodstream and urinary tract infections in critically ill or immunocompromised patients. Its ability to persist in healthcare environments, acquire and disseminate antibiotic resistance genes, and colonize the gastrointestinal tract has driven its emergence as one of the most important nosocomial pathogens worldwide (1, 2). Vancomycin-resistant *E. faecium* (VR*Efm*) strains present significant therapeutic challenges, primarily due to limited treatment options and their potential for rapid clonal spread within healthcare settings (3). In recent years, the epidemiology of VR*Efm* in Europe has shown largely stable, country-dependent trends, with few nations reporting increases (e.g., Estonia, Spain and Italy) and others, such as Germany and Ireland, noting declines in invasive isolates (www.ecdc.europa.eu). In Germany, these overall trends are complemented by shifting patterns in *van* genotypes (*vanA* and *vanB*) and clonal lineages observed across time and geography (4).

Whole-genome sequencing (WGS) has become the gold standard for the detailed characterization and typing of VR*Efm*, with cgMLST as a key WGS-based approach. CgMLST outperforms traditional methods such as PFGE and MLST by providing enhanced resolution through the analysis of 1,423 core genes (5). Over the past decade, recommendations for setting the allelic threshold to identify outbreak-related clusters using cgMLST have shifted markedly - from early cutoffs of up to 20 alleles to more recent proposals of ≤3 allelic differences (5-8).

Since 2016, the ST117/CT71 lineage has emerged as the dominant clone among invasive VR*Efm* isolates in Germany (9, 10). WGS analysis from the German National Reference Center (NRC) for Staphylococci and Enterococci has revealed its widespread dissemination over the past years, peaking around 2019, followed by a notable decline. However, when applying cgMLST for strain typing, isolates from this lineage without an apparent epidemiological link often appear highly related, which can complicate outbreak detection. Recent studies have shown that integrating split k-mer–based analyses can overcome this limitation by providing markedly higher discriminatory resolution for closely related *E. faecium* isolates (11, 12).

This study aims to assess the efficacy of cgMLST and split k-mer analysis (SKA), in distinguishing ST117/CT71 isolates, while also exploring the potential of transposon reconstruction to further enhance genomic resolution. We investigate whether limitations in strain discrimination by cgMLST are specific to certain clonal lineages or reflect a broader feature of the *E. faecium* population structure. Finally, we analyze clinical outbreak events to identify allelic differences within confined settings, with the aim of establishing a foundation for adjusting the threshold for cgMLST analysis.

## Results

### Epidemiology of invasive *E. faecium* in Germany from 2015 – 2024

All *E. faecium* isolates associated with bloodstream infections (BSI) submitted to the NRC have been routinely typed by cgMLST, allowing us to track epidemiological trends in strain prevalence over time in Germany. The distribution of invasive VR*Efm* observed from 2015 to 2024 (n=1,142) revealed significant changes in the frequencies of specific STs and CTs (Figure 1). VR*Efm* belonging to ST117 exhibited a marked increase in prevalence, peaking in 2019 and maintaining its dominance until 2024. Central to this trend is ST117/CT71, which experienced a sharp increase up to 2019, followed by a decline in subsequent years (Figure 1). ST80 followed a gradual yet consistent upward trajectory over the same period. In addition, ST1299, which had previously shown only limited expansion, exhibited a marked increase in 2024, with 31 isolates identified, 24 of which belonged to CT3109.

**Figure 1.**
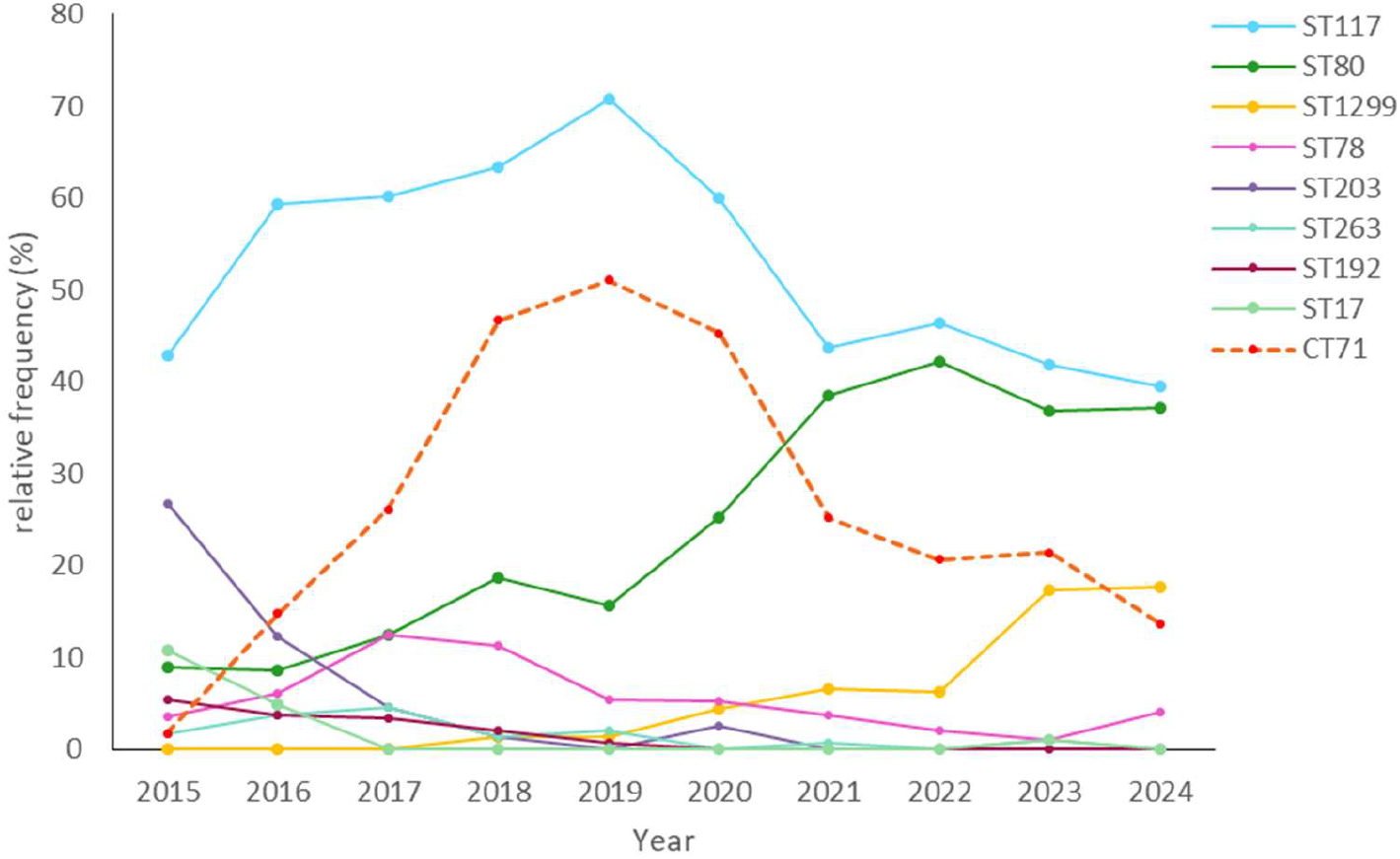
Temporal trends in the distribution of dominant STs of invasive VR*Efm* in Germany from 2015 to 2024 (n=1,143). Each ST is represented by a distinct colored line, with CT71 highlighted by a red dashed line.

### Molecular Typing and Transposon Analysis of invasive *E. faecium* ST117/CT71

Between 2015 and 2024, as part of the NRC’s BSI surveillance, 297 *E. faecium* isolates were identified as ST117/CT71 (2015: n=1; 2016: n=9; 2017: n=15; 2018: n=61; 2019: n=57; 2020: n=51; 2021: n=34; 2022: n=22; 2023: n=22; 2024: n=25; Supplementary Table 1). All but three isolates were *vanB-*positive, corresponding with the respective resistance phenotype. Due to single nucleotide polymorphisms in one of the seven housekeeping genes, seven CT71 isolates were assigned to different STs. For simplicity, hereafter, we refer to these isolates as ST117/CT71. To investigate the genetic relationship among these ST117/CT71 isolates, a NJT was constructed based on cgMLST (Figure 2). By applying a clustering threshold of ≤15 and enabling the pairwise option, all ST117/CT71 *E. faecium* isolates – except for UW23514 – formed a single large cluster, despite being collected from different locations and time points (Figure 2, Supplementary Figure 1). To further distinguish the ST117/CT71 cluster, SKA was performed using a threshold of 7 SNPs (default). This high-resolution analysis identified 223 singletons and 30 distinct subclusters (Figure 2, Supplementary Figure 2). Notably, several SKA-defined clusters showed clear geographic or temporal coherence.

**Figure 2.**
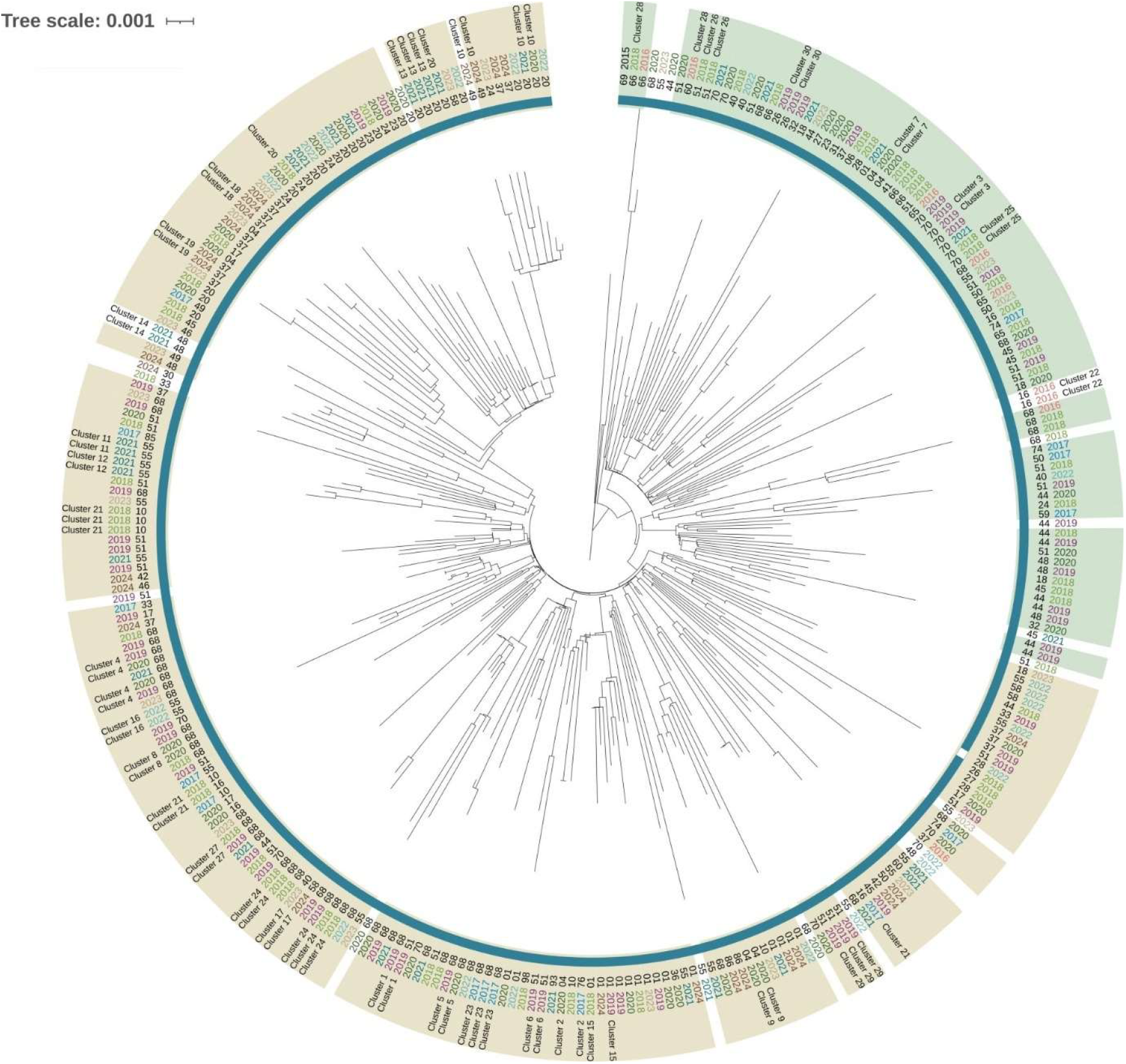
Bruker MBioSEQ Ridom Typer NJT of 297 invasive ST117/CT71 *E. faecium* samples based on 1,423 loci, constructed using the pairwise ignoring missing values option. Each branch tip represents an individual isolate labeled with a unique identifier (UW number). The inner dark blue circle denotes isolates grouped into a single cluster based on an allele threshold of 15. The second ring depicts the geographic location (2-digit zip code) of the submitting laboratory or hospital, and the third ring depicts the year of isolation, with distinct colors assigned to each year. The outermost circle specifies the subclusters identified by SKA analysis. The colored ranges define the identified transposon types (green = type 1, brown = type 2). Rare transposon types are not represented (white area).

In addition to cgMLST and SKA, we performed transposon analyses for all 297 ST117/CT71 *E. faecium* isolates. The analysis revealed two predominant Tn1549-like variants, designated type 1 (n=79) and type 2 (n=194) (Supplementary Table 1), both integrated into a gene of unknown function (HMPREF0351_10592). In addition, rare minor derivatives of both types 1 and 2 were observed in six isolates (data not shown). While both major variants show the insertion of two IS*Efa17* elements within the *vanB* operon (Figure 3), the Tn*1549*-like type 2 variant uniquely features an 18 bp duplication in the *vanS* gene and a 120 bp deletion in the *vanW* gene (Figure 3). Temporal analysis revealed a clear shift in the distribution of the two Tn*1549* variants over time: Type 1-associated *E. faecium* ST117/CT71 predominated between 2015 and 2018, whereas Type 2 emerged around 2017 and became dominant from 2019 onward, accounting for nearly all isolates by 2024 (Supplementary Figure 3). Consistent with these genetic and temporal distinctions, the two transposon structures were associated with distinct phylogenetic clusters (Figure 2). Validation through ONT sequencing and read mapping confirmed that both major variants were chromosomally integrated. In addition, analysis of the flanking chromosomal regions revealed only minor genetic differences. Notably, identical transposon structures were also detected in other clonal backgrounds, including ST117/CT36, ST262/CT1561, and ST80/CT1065 (data not shown).

**Figure 3.**
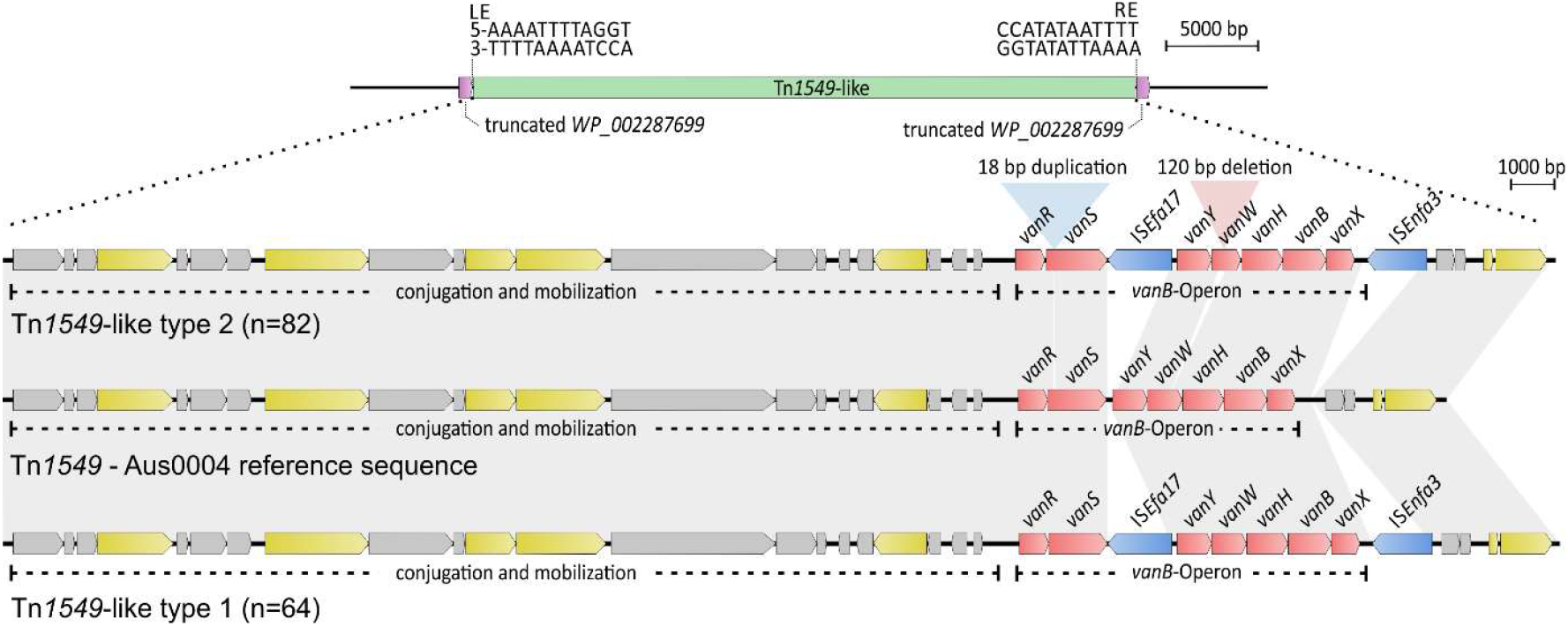
Schematic representation of Tn*1549*-like types 1 (bottom) and 2 (top) compared to the Tn*1549* reference structure of *E. faecium* Aus0004 (Accession number: CP003351). The upper panel shows the chromosomal integration site of Tn*1549*. In the detailed illustration of the Tn-structures, structural features – including genes involved in conjugation and mobilization as well as the *vanB* operon – are marked. Insertions and deletions relative to the reference sequence are indicated by white-shaded regions, whereas grey-shaded regions represent identical segments.

### Allelic profile of *E. faecium*: Key Findings from cgMLST Analysis

To establish a population-wide allelic profile of *E. faecium* circulating in Germany (VS*Efm* and VR*Efm*), we analyzed 2,505 isolates using cgMLST, which covered 48 different STs and 392 different CTs (Supplementary Table 2). These isolates, collected between 2014 and 2023, were linked to BSI, outbreak events, or the occurrence of rare antibiotic resistance mechanisms. Based on the allele profiles obtained from the cgMLST analysis, we calculated the allelic differences between all isolate pairs (Supplementary Table 3; available at zendod 10.5281/zenodo.23985109). In total, this included 3,136,260 isolate pairs with defined allele distances (after removing NA values). Across the studied *E. faecium* population, most pairwise comparisons ranged between 0 and 550 allelic differences (Figure 4a). When aggregating all allelic distances assigned to the category “same CT” across the dataset, isolate pairs exhibited a range from 0 to 37 allelic differences. These values formed a central peak at approximately 16 alleles (mean = 16.02) (Figure 4b and Supplementary Material 2). The allelic profile of CT71 isolates closely matched this pattern (mean = 16.55) (Figure 4b and Supplementary Material 2). Comprehensive graphical representations for individual CTs – including CT71, CT929, CT2505, CT1470, CT1065 and CT469 – are provided in Supplementary Material 1.

**Figure 4.**
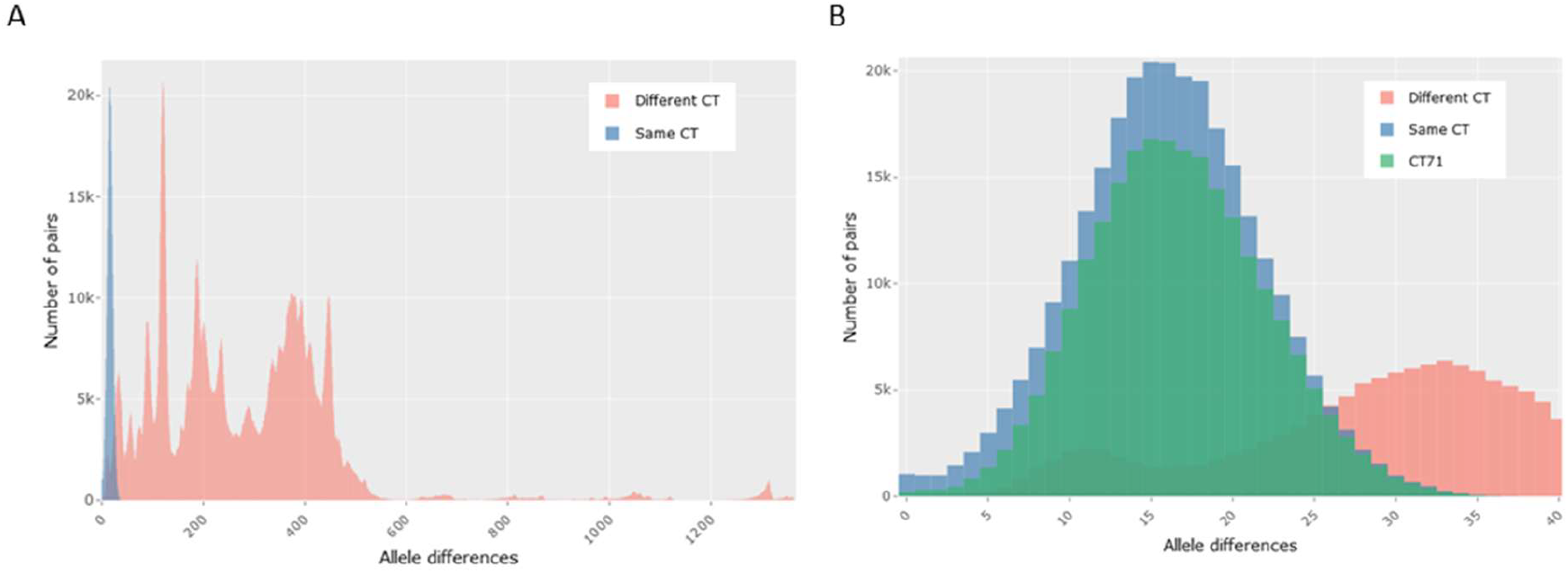
Overlaid Histograms showing the distribution of cgMLST allelic differences between all *E. faecium* isolate pairs (*n* = 2,505). (A) Comparisons within the same CT are shown in blue, whereas comparisons between different CTs are shown in red. (B) Zoomed-in view with the allele distribution of the CT71 isolates highlighted in green.

When comparing the allele distances among the dominant CTs of *E. faecium* circulating in Germany between 2014 and 2023 (Figure 5), isolates belonging to CT71 (highlighted in blue) showed comparable allelic differences than those of other cluster types (Q2/median: CT71=16, CT929=9, CT1065=15, CT1470=16, CT2505=16, CT469=13). Notably, CT929 showed a median allelic difference of fewer than 10 alleles (Q2/median = 9) (Figure 5). To assess whether allelic distance distributions differed between CTs, we used a non-parametric Mann–Whitney U test, as the data were discrete and not normally distributed. Although p-values were extremely low – indicating statistically significant differences between CTs – the effect sizes were modest, suggesting that these differences are of limited practical relevance. Furthermore, given the large sample size, even minor differences may reach statistical significance. Full analyses as well as statistical analysis are provided in Supplementary Material 1 and 2, respectively.

**Figure 5.**
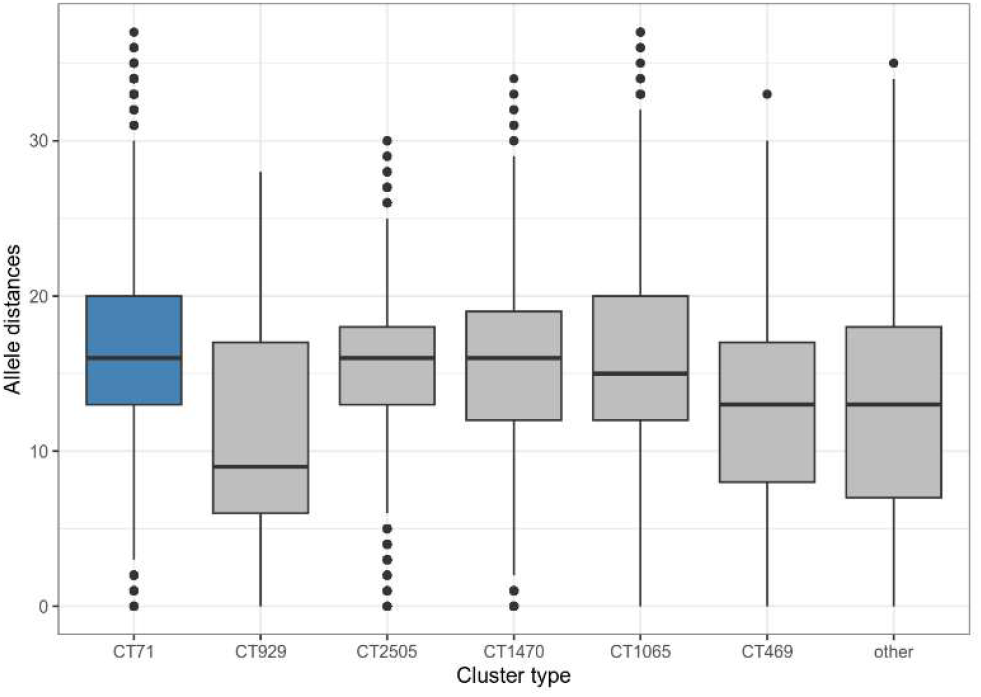
Boxplots of all allele distances for a given cluster type. Boxplots show the range of pairwise allelic differences among isolates belonging to the selected CTs, with CT71 highlighted in blue. Outliers are indicated by individual points.

### Use of cgMLST for the analysis of *E. faecium* outbreaks in hospital settings

Between 2019 and 2024, the NRC for Staphylococci and Enterococci was contacted 161 times to support in outbreak investigations. To better understand the allelic differences within hospital-associated outbreak clusters, 96 single hospital outbreaks were analyzed (2019: n = 9; 2020: n = 26; 2021: n = 26; 2022: n = 12; 2023: n = 12; 2024: n = 11), comprising 138 distinct clusters (2019: n = 14; 2020: n = 40; 2021: n = 35; 2022: n = 16; 2023: n = 15; 2024: n = 24) (Supplementary Table 4). The contributing laboratories and hospitals were distributed across Germany without regional clustering (data not shown). The most frequently identified STs were ST117 (n = 62), ST80 (n = 46), and ST1299 (n = 12). Among the CTs, CT71 (ST117, n = 37) and CT1065 (ST80, n = 16) were the most common. On average, each outbreak comprised approximately four isolates, with the largest and smallest outbreaks including 17 and two isolates, respectively. Most of the allele distances within the outbreak clusters were concentrated at the lower end of the distribution (Figure 6). The histogram displays a pronounced peak at 0 (*n*=219) and 1 (*n*=82) allelic differences, followed by a steep decline in counts with increasing allele distance. Distances greater than three alleles occurred rarely. The overlaid boxplot indicates a median of one allelic difference, with the interquartile range extending from 0 (Q1) to 2 (Q3). The percentile values were distributed as follows: 5^th^–25^th^ percentile = 0, 50^th^ percentile (median) = 1, 75^th^ percentile = 2, 80^th^ percentile = 3, 85^th^ percentile = 5, 90^th^ percentile = 9, and 95^th^ percentile = 12. Outliers corresponding to the upper end of the distribution were also observed (Figure 6).

**Figure 6.**
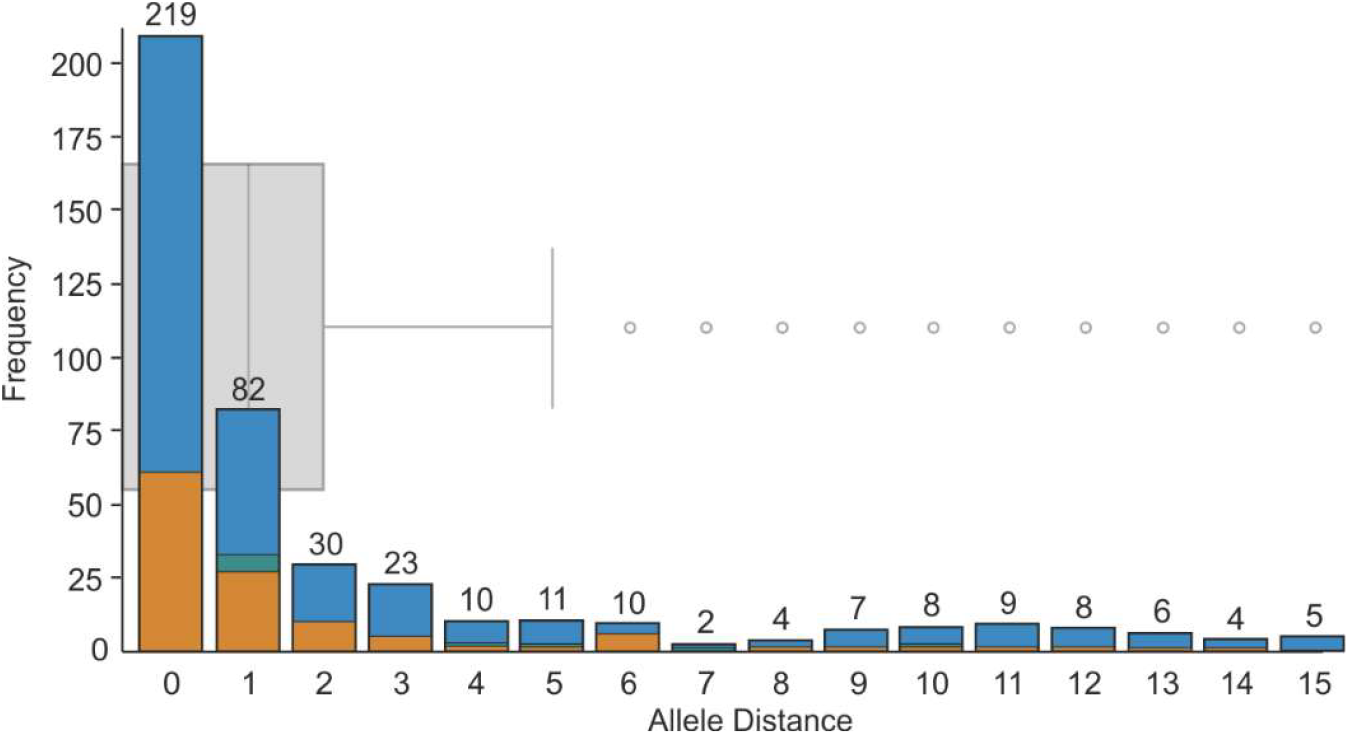
Distribution of allele distances within 138 outbreak clusters (n = 96 outbreaks, 2019–2024). The stacked histogram depicts the counts of pairwise allelic differences (ST80/CT1065 (green), ST117/CT71 (orange), other clonal lineages (blue)). The overlaid box plot (grey) summarizes the median, interquartile range, and the outliers.

## Discussion

Recent studies have shown that *E. faecium* ST117/CT71 cannot be reliably resolved using conventional typing methods such as cgMLST (9, 12). Using this lineage as a model, we assessed the resolution limits of current typing approaches and, supported by population-wide data, examined whether reduced genomic diversity within the core-genome represents a broader feature across dominant *E. faecium* lineages. Our analyses show that cgMLST fails to sufficiently resolve ST117/CT71 isolates and that high-resolution strategies such as SKA are required to achieve finer genomic discrimination. Moreover, population-wide analyses confirmed that reduced core-genome diversity is common among dominant *E. faecium* lineages and not specific to CT71, highlighting the need to critically reassess current thresholds and methodologies for genomic typing.

Longitudinal genomic surveillance conducted at the NRC reveals that the population structure of invasive *E. faecium* in Germany is characterized by successive clonal shifts. A particularly striking example of this dynamic is the hospital-adapted lineage ST117/CT71, which dominated between 2016 and 2021 (9, 10, 13). Similar temporal patterns were observed for other lineages by our NRC, albeit to varying extents (e.g. ST80/CT1065, ST80/CT1470). Notably, since 2023, we have detected a pronounced rise of isolates belonging to ST1299/CT3109 – a clonal lineage not detected by the NRC prior to 2021. This emergence aligns with recent reports documenting the rapid dissemination of *E. faecium* ST1299/*vanA* across Southern and South-Western Germany and Austria (14, 15). Likewise, patterns of adaptive turnover have been observed in neighboring countries, including Denmark, the Netherlands, and the United Kingdom (16-18). Such dynamics are likely shaped by the combined effects of antibiotic selection pressure, fitness costs associated with plasmid carriage, ecological competition, and adaptive genomic evolution within the hospital niche (19).

In the genomic epidemiology of *E. faecium*, cgMLST has become the gold standard for surveillance and outbreak detection. Unlike traditional MLST, cgMLST analyzes more than 1,400 core genes, thereby overcoming the limited discriminatory power of conventional typing schemes (20). However, in our analysis of 297 invasive ST117/CT71 *E. faecium* isolates collected between 2015 and 2024, almost all samples clustered within a single cgMLST cluster (≤15 allelic differences). Epidemiologically, such uniform clustering across different hospitals, regions, and years is highly implausible, suggesting that cgMLST underestimate genetic differences. Similar findings were reported by Falgenhauer *et al*. (2019), who identified a near-ubiquitous ST117/CT71 clone circulating in multiple geographically separated institutions of the Rhine-Main area of Germany (9). Together, these observations indicate that a relatively stable core-genome profile of ST117/CT71 isolates severely limit the capacity of cgMLST to distinguish epidemiologically unrelated strains. This limitation is particularly pronounced for dominant clonal lineages with a high incidence. The limited discriminatory power of cgMLST for dominant *E. faecium* lineages has likewise been demonstrated by a recent study, showing that cgMLST often failed to resolve suspected transmission chains and outbreaks involving hospital-adapted lineages such as ST79, ST80, and ST117 (12). Given the insufficient resolution of cgMLST, we implemented SKA (21) as a complementary high-resolution approach. The superior discriminatory power of SKA has been demonstrated by Maechler *et al*. (2023), who emphasized its ability to capture additional genomic variation, including plasmid-associated content. Indeed, this method resolved the large cgMLST cluster into several smaller subclusters that correlated more closely with epidemiological data. However, as the reference-free SKA approach was not designed for large-scale population genomics, it should be applied in a hierarchical workflow (11).

Previous studies have emphasized that incorporating analyses of mobile genetic elements (MGEs) can enhance the discriminatory power of genomic typing and outbreak investigation (22-26). Accordingly, we performed a comprehensive transposon analysis of all 297 *E. faecium* ST117/CT71 isolates, identifying two *Tn1549*-like variants, both integrated at the same chromosomal locus corresponding to a gene of unknown function (HMPREF0351_10592). This aligns with previous reports from Germany, where identical insertion of *vanB*-carrying *Tn1549* elements into HMPREF0351_10592 was observed for the *E. faecium* lineages ST117 and ST192 (9, 27). Our findings also align with the work of Howden *et al*., who demonstrated that Tn*1549* insertion is not random: among 36 VR*Efm* isolates they identified a restricted repertoire of chromosomal integration sites. Although seven distinct insertion sites were detected, transposons of identical sequence consistently integrated into the same chromosomal locus (24). Taken together, the recurrence of this integration hotspot across distinct *E. faecium* backgrounds, combined with the limited set of permissible insertion sites described previously, suggests that specific chromosomal environments promote preferred acquisition and long-term stable integration of Tn*1549*.

Interestingly, both Tn-variants were each linked to a distinct phylogenetic clade and occurred during different time periods, revealing that the population previously defined as ST117/CT71 actually comprises two genetically distinct sublineages circulating at different times in Germany. This observation suggests that *vanB* was either acquired independently at least twice during the evolution of this lineage – through repeated exposure to locally circulating mobile elements – or that Type 2 evolved directly from Type 1 *via* minor structural modifications of an already integrated Tn*1549* element. The latter scenario is supported by the near-identical flanking chromosomal regions, although previous work by Bender *et al*. indicated that larger chromosomal fragments carrying *vanB*- associated Tn*1549* can also be exchanged between closely related isolates (27).

As *E. faecium* ST117/CT71 has frequently been described as highly clonal based on cgMLST, we sought to determine whether pairwise allelic distances within this lineage were smaller than those observed across the broader *E. faecium* population or compared with other dominant lineages. To this end, we performed a population-wide cgMLST analysis encompassing 2,505 *E. faecium* isolates. The allelic distance distribution observed for ST117/CT71 closely mirrored the *intra*-CT distribution of all other lineages, indicating that CT71 is not more clonal at the cgMLST level than other hospital-adapted clonal lineages of the A1 clade. Thus, our findings refute the initial assumption that CT71 exhibits unusually high clonal stability. This conclusion is further supported by comparisons with other dominant CTs, where CT71 displayed a median allelic distance similar to or slightly higher than CT1065, CT1470, and CT2505. Because our dataset represents a broad background population sampled from across Germany over a ten-year period, it is expected to include mostly unrelated isolates. The population-wide allelic peak at approximately 15 alleles difference therefore likely reflects the intrinsic genomic diversity of circulating *E. faecium* lineages rather than a signal of recent clonal spread. Low levels of nucleotide diversity across core genes within clade A1 *E. faecium* have previously been demonstrated by van Hal *et al*., who proposed that clade A1 likely originated from the expansion of a single ancestral clone followed by decades of diversification (28). Our observation indicates that even epidemiologically unrelated isolates from the general background population can display cgMLST distances well below the conventional outbreak threshold of ≤20 (5). Consequently, applying high cgMLST thresholds for cluster definition in the absence of supporting epidemiological data may result in misleading inferences of transmission, particularly for dominant lineages that are highly prevalent.

Building on the observation that the general background population already centers around ∼15 cgMLST alleles difference, we next examined whether hospital outbreak clusters display a distinctly tighter genetic signal. The distribution of allelic distances in these outbreaks was highly compressed, with most isolate pairs exhibiting only minimal allelic differences, while larger distances were observed only rarely. Notably, this tight clustering reflects geographically and temporally confined outbreaks, with the limitation that only pairwise allelic differences, not distances derived from distance matrices, were considered. Although such findings might argue for implementing highly stringent thresholds – as suggested and used in previous outbreak studies (6, 7) – these should be applied with caution. While tighter cutoffs increase specificity, they risk fragmenting genuine transmission networks when sampling is incomplete, within-host diversity occurs, mutations accumulate during prolonged outbreaks, or when multicenter outbreaks emerge. This, in turn, may lead to an overall reduced awareness and vigilance in clinical settings. Importantly, sequencing at the RKI is performed using a validated and highly standardized laboratory and bioinformatic workflow, thereby minimizing technical variability. Such consistency cannot be assumed across all medical, public-health, or research institutions; differences in sequencing platforms, laboratory workflows, or versions of bioinformatic tools and pipelines are well known to introduce small allelic artefacts, thereby inflating genomic distances (29, 30).

Together, the population-level and outbreak data delineate a clear and practical distinction: while isolates within nationwide circulating lineages typically differ by around 15 alleles, epidemiologically linked isolates cluster almost exclusively within 0-3 alleles. To better balance sensitivity and specificity in outbreak detection – while avoiding the fragmentation of genuine clusters – our findings support adopting a cgMLST alert threshold of ≤ 9 as a pragmatic starting point for cluster detection. This threshold corresponds to the 90^th^ percentile of allelic distances observed within investigated outbreaks, thereby capturing the vast majority of outbreak-related links while remaining sufficiently below the ∼15-allele background signal of circulating lineages to maintain discriminatory power. Such an approach should then be followed by high-resolution confirmation using SKA analyses, as also proposed by Higgs *et al*. (11), with genomic clustering signals interpreted in conjunction with epidemiological data. Importantly, this proposed threshold is particularly relevant for laboratories conducting regional or supra-regional surveillance, where detailed epidemiological information is often limited. We therefore encourage future studies to further examine and challenge both the threshold and the stepwise analytical framework using well-characterized outbreak examples.

Our findings highlight that the limited core-genome diversity in clade A1 *E. faecium* is a population-wide characteristic of hospital-adapted lineages rather than a lineage-specific feature of ST117/CT71, fundamentally restricting the resolving power of cgMLST. To better balance sensitivity and specificity in outbreak detection, we propose a data-driven cgMLST alert threshold of ≤9 alleles, which captures most outbreak-related links while remaining below the ∼15-allele background of nationally circulating lineages. High-resolution methods such as SKA should be applied for detailed outbreak analysis. In addition, transposon analyses identified two distinct ST117/CT71 variants that circulated across Germany during the period of high VRE prevalence, providing further insight into the subtle microevolutionary dynamics that can occur even within seemingly homogeneous dominant lineages.

## Material and Methods

### Routine Testing and Molecular Typing of *Enterococcus* spp. Isolates at the NRC

This retrospective study analyzed *Enterococcus* spp. isolates submitted to the NRC between 2014 and 2024. The isolates originated from hospitals and diagnostic microbiology laboratories across Germany and were subjected to a standardized workflow comprising antimicrobial susceptibility testing, species-level identification, and determination of vancomycin resistance determinants (*van* genes). Testing procedures and result interpretation followed standardized protocols and clinical breakpoints established by EUCAST. Species identification and *van* gene detection were performed using PCR. In the case of isolates from blood cultures, suspected outbreaks, or strains displaying uncommon antibiotic resistance, WGS was performed on an Illumina platform. For reference, the general procedures for DNA extraction and sequencing are briefly summarized: For bacterial cultivation, Brain Heart Infusion medium was used, and genomic DNA was extracted from overnight cultures using the DNeasy Blood & Tissue Kit (Qiagen, Hilden, Germany). DNA concentrations were measured using the Qubit dsDNA HS Assay Kit and Qubit 4 Fluorometer (Thermo Fisher Scientific, Karlsruhe, Germany). Sequencing libraries were prepared using the Nextera XT DNA Library Preparation Kit (Illumina, San Diego, CA, USA) and paired-end sequencing was performed on an Illumina platform (Illumina MiSeq, Illumina HiSeq 1500, NextSeq 550 or NextSeq 2000) with a minimum coverage of 70×.

### Molecular Typing and Genomic Characterization of ST117/CT71 *E. faecium* isolates

#### Whole Genome Sequencing, Quality Control, and Genome Reconstruction

Between 2014 and 2024, a total of 336 ST117/CT71 isolates were sequenced at the NRC. Available raw sequencing data were reassessed for quality using FQStat (v0.0.12) (31) and Bruker MBioSEQ Ridom Typer. Reads passing quality checks (*n*=297 isolates) were subjected to *de novo* assembly with SPAdes (v3.15.4) (32), applying default parameters, and enabling the ‘careful’ mode (Supplementary Table 1).

#### Phylogenetic Analysis

MLST and cgMLST analyses were performed using *de novo* assembled contigs in Bruker MBioSEQ Ridom Typer. The cgMLST task template described by de Been *et al*. was applied (5), and cgMLST clusters were defined using a pairwise allelic difference of ≤15. A neighbor-joining tree (NJT) was constructed from the cgMLST data, with pairwise missing values excluded, and visualized using iTOL (v7) (33). Epidemiological metadata were integrated into the NJT, whereby the geographic origin was assigned based on the location of the submitting clinical laboratory (*via* partial 2-digit zip code) (Supplementary Table 1). To refine phylogenetic resolution, a reference-free SKA (https://github.com/simonrharris/SKA) was performed (34). K-mer files (k=15) were generated from the raw sequencing data using the *fastq* subcommand with default parameters. Pairwise distances were calculated and subjected to single linkage clustering. Clusters were defined using a threshold of seven SNPs (default settings). For illustration purposes, we used RAWGraphs (35).

#### Transposon Reconstruction

To identify *vanB*-carrying contigs, *de novo* assembled contigs were aligned to the *vanB operon* sequence of *E. faecium* Aus0004 (CP003341, positions 2,838,182–2,846,134) using Geneious Prime 2025.0.2. When mismatches occurred at overlapping contig ends that did not align to the reference sequence, we examined the regions for inverted repeat motifs using a customized database of annotated IS elements and their corresponding IRL/IRR signatures. If complementary inverted repeats were identified at both contig ends, the appropriate IS element was manually inserted between the two repeats to reconstruct the missing junction. Additionally, wherever insertions or deletions were detected, the respective structural changes were incorporated accordingly. The structural integrity of the reconstructed transposon sequence was then verified by aligning trimmed Illumina reads. For validation, long-read sequencing of selected isolates was performed using Oxford Nanopore Technology. Bacterial isolates were cultivated in LB medium for 4 h, after which 2 mL of culture was centrifuged at 16,000 rpm for 2 min. The resulting pellet was subjected to high-molecular-weight DNA extraction using the Wizard® HMW DNA Extraction Kit (Promega, Fitchburg, USA) on an EpMotion 5075tc automated pipetting system (Eppendorf, Hamburg, Germany). Sequencing libraries were prepared using the SQK-NBD114-24 Kit and sequenced on a GridION device equipped with a FLO-MIN114 R10.4.1 flow cell (Oxford Nanopore Technologies, Oxford, UK). Base calling was performed in super-accurate mode with the following parameters: modified base calling off, trim barcodes on, mid-read barcode filtering off, and a minimum Q score of 10. The data quality was monitored using MinKNOW (v24.11.8). Hybrid *de novo* assembly was subsequently performed by combining long (ONT) and short (Illumina) reads using Unicycler (v0.5.0) in conservative mode (36).

### Population wide allelic profile of *E. faecium*

In order to establish a population-wide allelic profile of *E. faecium* circulating in Germany, a total of 2,505 isolates collected between 2015 and 2023 were analyzed using cgMLST (Supplementary Table 2). For all isolates, *de novo* assembled contig-files were available at the time of analysis (see section Routine Testing and Molecular Typing of Enterococcus spp. Isolates at the NRC). Their quality was subsequently re-evaluated using Bruker MBioSEQ Ridom Typer (Ridom; Münster, Germany, client version 10.5.4). Pairwise allele distances were calculated for all pairs of the 2,505 isolates based on their allelic profiles. The distance matrix that was compiled into an Excel spreadsheet (Supplementary Table 3; available at zenodo: 10.5281/zenodo.23985109). Subsequent analyses were performed in R (v4.4.3) using R Markdown (v2.29) for documentation and report generation. Each record represented a pair of isolates with associated allele distance, ST, and CT information. Allele distance histograms were created for all isolates, with additional groupings by ST or CT. Further analyses compared allele distances among isolates with identical versus different ST or CT. All visualizations were produced using ggplot2 (v3.4.3), with interactive versions made available through plotly (v4.11.0). Descriptive statistics, including the mean, median, standard deviation, quartiles, minimum, and maximum, were calculated for selected CTs using functions from the stats package (v4.4.3). To illustrate distributions, histograms with superimposed normal density curves and boxplots were created. Given that allele distances are discrete and do not follow a normal distribution, non-parametric tests were employed. Specifically, Mann–Whitney U tests (also known as Wilcoxon rank-sum tests for independent samples) were conducted to compare the allele distance distributions of CT71 with those of other selected CTs (929, 2505, 1470, 1065, 469). The significance level was set at α = 0.05. Effect sizes were determined to evaluate the magnitude of differences between groups. The complete analysis process, including plot creation, is accessible as an R Markdown script for reproducibility if requested. The complete report can be found in the Supplementary Material 1 and Supplementary Material 2.

### Characterization of allele distance distributions in outbreak-related clusters

To systematically evaluate genomic relatedness in potential outbreak events, 96 single hospital outbreaks (outbreaks occurring within a single hospital) were included in the analysis, encompassing 138 distinct cgMLST clusters and 577 *E. faecium* isolates collected from 2019 to 2024 (Supplementary Table 4). Clusters were defined by an alert threshold of ≤15 allelic differences and included when they occurred within a temporally restricted period at a single medical institution. Only clusters with a single CT were analyzed, and duplicate patient samples were omitted. Distances were derived from the MST. Allelic distance distributions within outbreak clusters were extracted from the dataset, summarized using frequency tables, and further described using statistical measures, including the median, interquartile range, and percentiles. The results were visualized as histograms with overlaid boxplots using Microsoft Excel and CorelDraw X6 (v16.4.1.1281) software.

## Data availability

The genome sequencing data generated in this study have been deposited in the European Nucleotide Archive (ENA) under the accession number PRJEB104440. Supplementary table 3 is available at https://doi.org/10.5281/zenodo.23985109.

## Acknowledgement

We would like to thank the team at the Sequencing Core Facility of the Genome Competence Centre, Robert Koch Institute, especially Aleksandar Radonic and Tanja Pilz, for their excellent support with Illumina and ONT sequencing of our samples as well as Sandra Kaiser and Simon Tausch for their bioinformatic support. We also thank all laboratories that provided us with strains, as well as Carola Fleige and Christine von Bornstädt for their dedicated laboratory support. This study was supported by a grant from The Ministry for Health to R.E.W. under the call for “Establishing an Integrated Genomic Surveillance for Germany” (2022 – 2025). We are grateful to all members of the VRE BSI Study Group submitting isolates to the NRC for further analyses, especially Institut für Medizinische Mikrobiologie und Virologie in Dresden, Institut für Medizinische Mikrobiologie in Göttingen, Klinische Mikrobiologie und Krankenhaushygiene in Oberhausen, Uniklinik RWTH Aachen, Klinikum Mannheim GmbH – Universitätsklinikum, Medizinische Universität Lausitz - Carl Thiem in Cottbus, Friedrich Loeffler Institut in Greifswald, Medizinische Universitätslaboratorien in Jena, Medizinisches Versorgungszentrum in Ingelheim, MEDILYS Laborgesellschaft mbH in Hamburg, Schön Klinik Rendsburg, SYNLAB in Augsburg, Labor Berlin - Charite Vivantes GmbH, Institut für Medizinische Mikrobiologie und Immunologie in Köln, LADR Laborzentrum Nord in Flintbek, UKD Universitätsklinikum Düsseldorf, Laborarztpraxis Osnabrück in Georgsmarienhütte, Labor Dr. von Froreich GmbH in Hamburg, ELBLAB GmbH in Meißen, Institut für Med. Mikrobiologie und Hygiene in Homburg, KLG Krankenhaus-Labor-Gesellschaft mbH in Frankfurt, Medlabwahl GmbH & Co. KG in Lüdenscheid, Universitätsklinikum Magdeburg A.ö.R., Institut für Klinische Mikrobiologie und Hygiene in Regensburg, Institut für Med. Mikrobiologie und Hygiene in Tübingen, Klinikum Ingolstadt. We acknowledge Zenodo for hosting supplementary material.

